# Pericytes promote more angiogenesis and vasculogenesis than mesenchymal stromal cells in microfluidic chips

**DOI:** 10.1101/2023.11.02.565266

**Authors:** Julian Gonzalez-Rubio, Hannah Kubiza, Hiltrud Koenigs-Werner, Stefan Jockenhoevel, Christian G. Cornelissen, Anja Lena Thiebes

## Abstract

The pericyte is a key player in vascularization, protecting endothelial cells from external harm and promoting formation of new vessels and connections when necessary. However, pericytic identity and its relation with other cell types, such as the mesenchymal stromal/stem cells, is highly debated. To compare the behaviour of pericytes and unselected stromal cells in vascularization, we used multichannel microfluidic chips to replicate the two processes of vessel formation: vasculogenesis, the *de novo* vessel formation, and angiogenesis, the sprouting of new vessels from existing ones. In angiogenesis, pericytes promote significantly more and longer sproutings than stromal cells. In vasculogenesis, stromal cells promote the formation of structures ressembling the embryonic capillary plexus, whereas pericytes wrap around the endothelial cells, arresting their division and forcing them into tubulogenesis. Whole-transcriptome sequencing confirms an upregulation of pro-vascularization and cytostatic genes in co-cultures of pericytes and endothelial cells; while stromal cells strongly stimulate mitosis and organelle biogenesis pathways and decrease the release of pro-inflammatory cytokines. In this study, we offer new insights into the pericyte-endothelial cell relation and the mesenchymal stromal cell elusive identity, relevant in both vascular biology and tissue engineering.

## 1. Introduction

The circulatory system is the interconnected and hierarchical network of vessels that irrigate most human tissues, nourishing the cells with nutrients and oxygen, removing waste products, and serving as a connection for the endocrine and immune systems to swiftly reach any part of the body (Yang *et al*., 2020). All vessels are lined with endothelial cells. In the capillaries, the smallest branch of the circulation, endothelial cells are lined externally by a mural cell type known as pericyte, which supports its homeostasis, protects it from stromal debris by phagocytosis, and regulates the blood flow (Zhao and Chappell, 2019).

For tissue engineering, the size of non-vascularized bioengineered organs is limited to a few hundred micrometers (Chandra and Atala, 2019). Any thicker tissue without irrigation of a blood-mimicking fluid will eventually lead to cell death due to the lack of oxygen and nutrient diffusion to the innermost parts, causing the formation of a necrotic core. However, achieving vascularization in *in vitro* systems is still a challenge due to the complexity of the process and the variety of cells involved (Dellaquila *et al*., 2021; Zhang and Radisic, 2020). The two main processes driving neovascularization are vasculogenesis, in which endothelial cells organize themselves into interconnected vessels, and angiogenesis, where new vessels originate from pre-existing ones.

Human adipose tissue is an easily obtainable biological source, which can be digested using collagenases to extract a heterogeneous mixture of cells, known as stromal vascular fraction (SVF) (Sun *et al*., 2019). The spindle-shaped cells produced by seeding the SVF on cell culture-treated plastic are often referred to as adipose-derived mesenchymal stromal/stem cells or adipose stromal cells (Bourin *et al*., 2013), in this study just termed stromal cells. They have been widely reported to have immunomodulatory properties (Melief *et al*., 2013) and *in vitro* multipotency (Brooks *et al*., 2019). Within this heterogeneous collection, we can find a wide variety of stromal cells such as fibroblasts, pre-adipocytes, and the previously mentioned pericytes (Sun *et al*., 2019). The Melanoma Cell Adhesion Molecule (MCAM), also known as Cluster of Differentiation (CD) 146, can act as a mural cell marker in bone marrow-derived stromal cells (Blocki *et al*., 2013) and has been proposed as a specific membrane protein for adipose-resident pericyte isolation (Wang *et al*., 2019). Importantly, MCAM is not exclusive to pericytes and can be expressed in other vascular-related cell types such as lymphocytes and endothelial cells, but these cells can be depleted due to their incapability or inefficiency to attach and divide on uncoated hydrophilic plastic surfaces (Wang *et al*., 2020). The identity of pericytes *in vivo* and, especially, *in vitro*, is however widely debated (Caplan, 2017; Maniyadath *et al*., 2023). So is their relationship with mesenchymal stem cells, their contribution to vascularization processes, and their use in cell therapies and tissue engineering (Galderisi, Peluso and Di Bernardo, 2022; Phinney, Hwa Lee and Boregowda, 2023; Viswanathan *et al*., 2019).

In recent years, microfluidic devices allowed the creation of organ-on-chip technologies, also known as microphysiological systems (Ingber, 2022; Leung *et al*., 2022). These platforms have been demonstrated as powerful tools for the study of tissue development, cell-cell and cell-matrix interaction, and vascularization (Alimperti *et al*., 2017; Vila Cuenca *et al*., 2021). Here, we use a microfluidic platform to mimic vasculogenic and angiogenic processes, comparing how pericytes and stromal cells interact with endothelial cells to form vessels and promote their growth. The present study is based on two hypotheses, (1) that CD146 serves as a reliable marker for the selection of pericytes from heterogenous adipose stromal cells, and (2) that pericytes can be cultured *in vitro* without losing the capacity to act as mural cells, promoting vessel formation better than their unselected counterpart and associating with the endothelial cells by a shared basement membrane. We investigate these assumptions at both the cellular and molecular levels using confocal and electron microscopy, as well as transcriptomic analysis.

## 2. Results and discussion

### 2.1. Microfluidic chip platform to study vessel formation via vasculogenesis and angiogenesis

To compare the vasculogenesis and angiogenesis processes (Figure 1A), and how pericytes and stromal cells act in either, we used a three-channel microfluidic chip (Figure 1B). To mimic vasculogenesis, endothelial cells and supporting cells – from now on a collective term for pericytes and stromal cells – are both added into the hydrogel in the middle channel, while for the angiogenesis only the supporting cells are added in the middle and one of the channels is coated with endothelial cells (Figure 1C). The pre-staining of the cells with lipophilic dyes confirms the deposition of the cells after gel injection. 24 hours after the preparation, endothelial cells are attached to the hydrogel-media interface in the angiogenesis condition, marking the beginning of angiogenesis (Figure 1C, right image).

**Figure 1.**
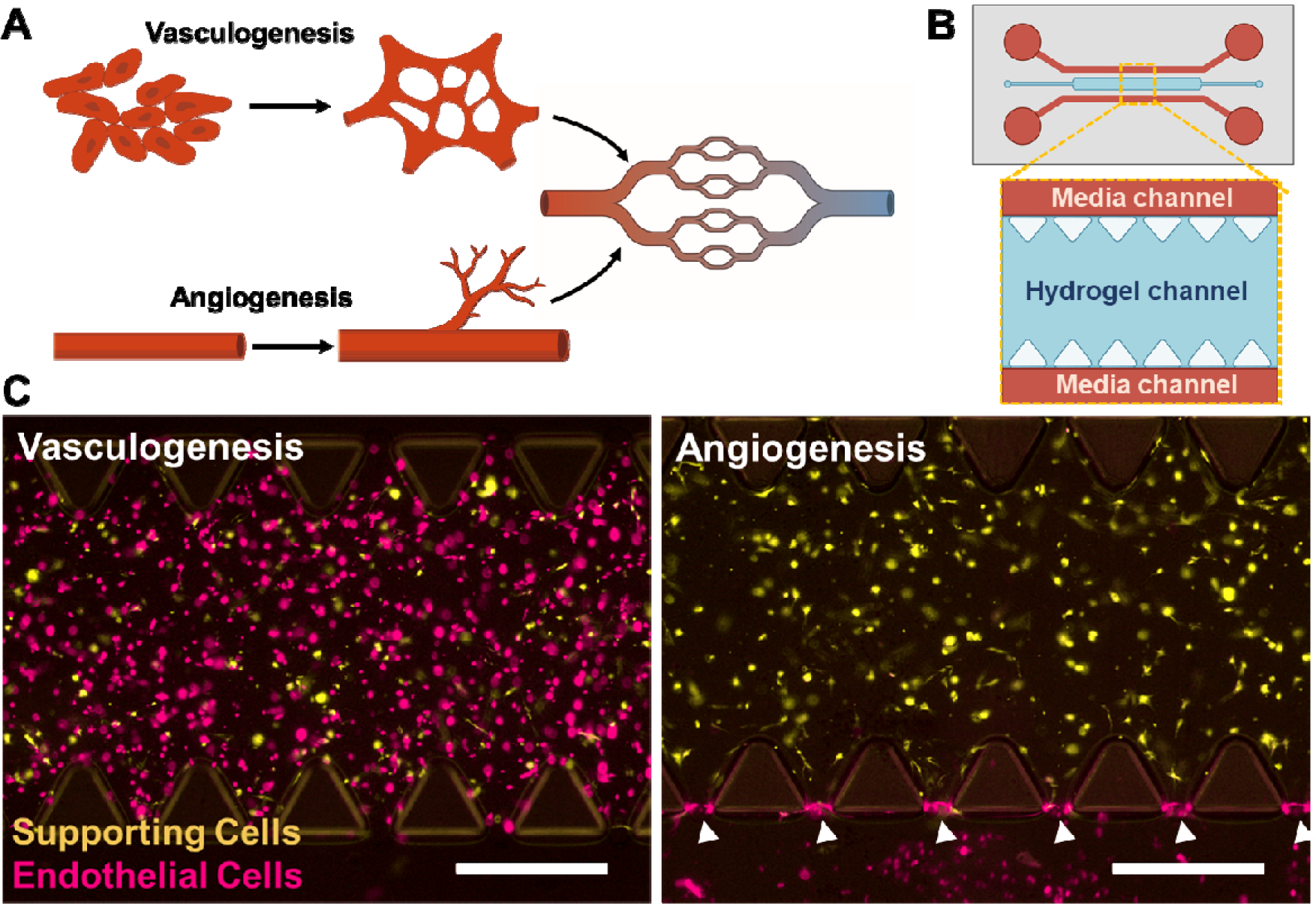
Replication of the two main neovascularization processes in microfluidic devices. **(A)** Main mechanisms of vessel formation in the human body. **(B)** Schematic drawing showing the internal hydrogel channel (blue) and the surrounding parallel media channels (red). **(C)** Fluorescence image showing endothelial cells (Vybrant DiD, magenta) and supporting cells (Vybrant DiO, yellow), which can be either pericytes or stromal cells, in two different set-ups to study vasculogenesis (left) and angiogenesis (right). White arrows mark the interface between hydrogel and media channel where the endothelial cells are attaching. Scale bars: 500 μm.

### 2.2. MCAM-enriched stromal cells act as pericytes *in vitro* and show enhanced pro-angiogenic potential

We separated pericytes from the heterogenous adipose SVF based on the expression of the mural cell marker MCAM/CD146 using immunomagnetic microbeads. Pericytes and normal unselected stromal cells were cultured in mesenchymal cell culture media and expanded for up to four passages. To investigate how these cell types compare to each other in promoting angiogenesis, three donors of each cell type were co-cultured with endothelial cells in microfluidic devices as described (Figure 1C, right). The co-cultures were maintained for ten days, fixed, and stained for the endothelial marker CD31, and for F-actin to mark the supporting cells. Confocal microscopy shows that pericytes are promoting more vessel sprouting than the stromal cells (Figure 2A). Statistically significant differences can be found in total vessel volume, length, and branching points (Figure 2B-D). Pericytes have been already reported to be highly pro-angiogenic supporting cells due to their close relation with endothelial cells *in vivo* (Blocki *et al*., 2013; Kosyakova *et al*., 2020; Yamazaki and Mukouyama, 2018). Interestingly, the increased angiogenesis is not correlated with any remarkable difference in vessel diameter (Figure 2E). Vessel lumen could not be confirmed or measured due to limitations in the confocal microscopy resolution. As can also be observed in the augmented immunofluorescence image, the pericytes tend to be significantly closer or even directly associated with the vessels, while the stromal cells remain homogenously distributed in the gel (Figures 2A and F). This interaction is expected from pericytes, which are known to wrap around blood vessels *in vivo* to support and regulate their functions (Yamazaki and Mukouyama, 2018). This way, pericytes can exert direct cell-cell pro-angiogenic stimulation (Caporali *et al*., 2017; Sweeney and Foldes, 2018). Apart from their relative position, pericytes do not show differences in the elongation with the stromal cells (Figure 5G). Overall, we can therefore confirm that the MCAM-selected pericytes maintain their pro-angiogenic activity and mural cell behavior *in vitro*.

**Figure 2.**
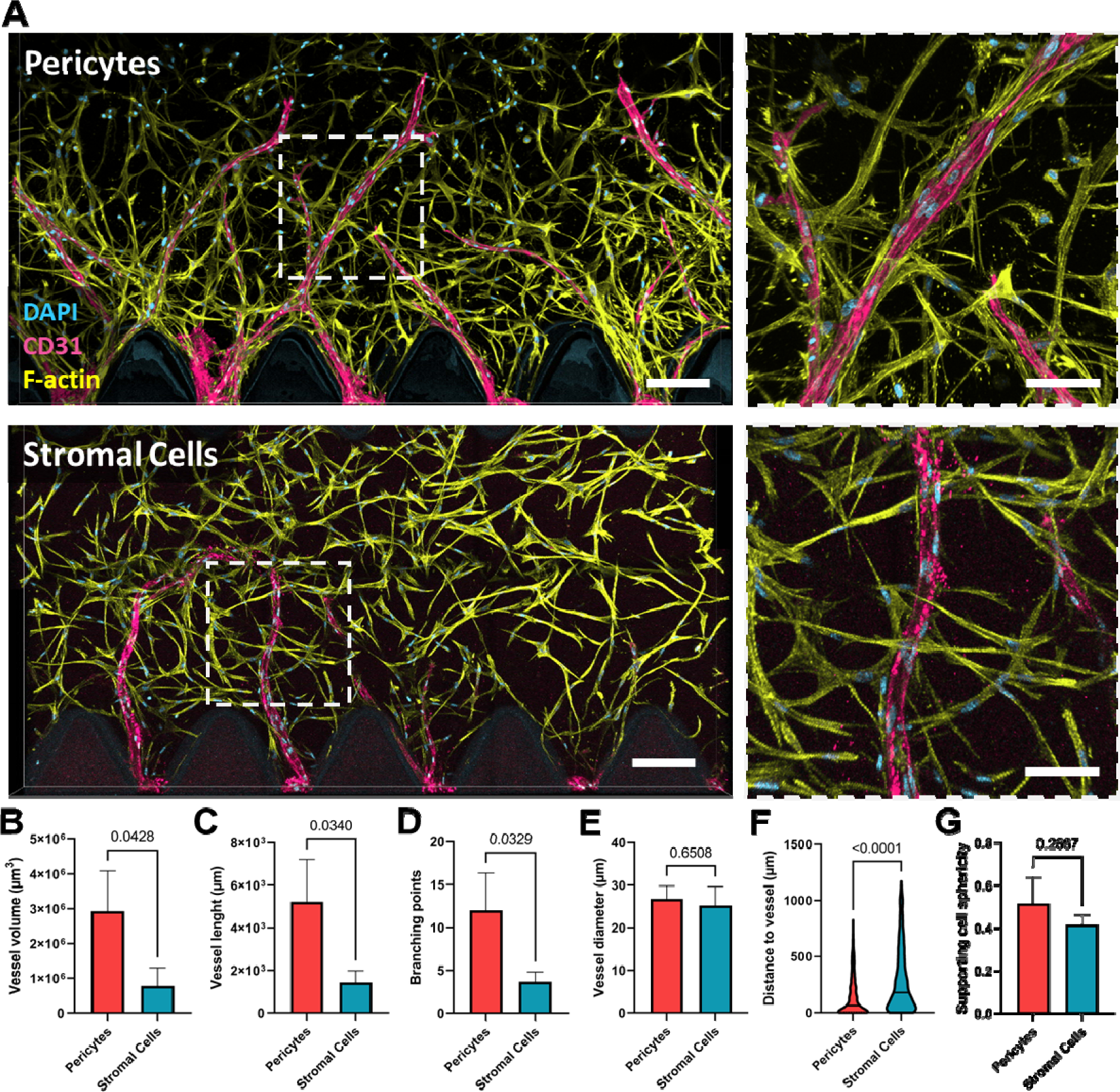
Comparison of the pro-angiogenic potential of pericytes and stromal cells. **(A)** Immunofluorescence images of the angiogenic sprouting within the microfluidic chips co-cultured with either pericytes or stromal cells, stained for nuclei (DAPI; cyan), CD31 showing the endothelial vessel-like structures (magenta) and the supporting cells actin cytoskeleton (Phalloidin; yellow). F-actin is not visible within the endothelial cells since it was subtracted from the picture during the image processing to increase clarity. Scale bar: 200 μm (left), 100 μm (right). **(B-E)** Comparison of vessels’ total volume, length, average diameter, and branching points between both co-culture conditions (n = 3, unpaired t-test, *p*<0.05 is considered significant). **(F)** Comparison of the distance between each type of supporting cell and the nearest vessel-like structure. The trunked violin plots depict summary statistics and the kernel density estimation to show the frequency distribution of each condition. The middle line represents the median (n = 3, Mann-Whitney test, *p*<0.05 is considered significant). **(G)** Comparison of supporting cell’s average sphericity (n = 3, unpaired t-test, p<0.05 is considered significant).

### 2.3. Pericytes promote vessel formation during vasculogenesis, while stromal cells stimulate endothelial cell proliferation

Immunofluorescence images of the microfluidic device comparing pericytes and the unselected stromal cells in vasculogenesis (Figure 1C, left) show clear differences between how both cell types support endothelial cell coalescence and elongation (Figure 3A). Pericytes promote the formation of long and thin tubular endothelial structures, while co-culture with stromal cells leads to the formation of bloated and amorphous structures. Quantification of the vessel-like structures shows a greater average area and volume in the gels co-cultured with stromal cells (Figure 3B and C). As shown by quantification of the total cell numbers, this is a consequence of a four-fold difference in the number of endothelial cells (Figure 3D). Stromal cells are significantly closer to the endothelial structures, although that closeness may relate to a lack of space due to the superlative endothelial structure size and cell number and not to a tight interaction between these cell types (Figure 3E). Altogether, pericytes promote the organization of endothelial cells into capillary-like structures while stromal cells stimulate their amorphous proliferation.

**Figure 3.**
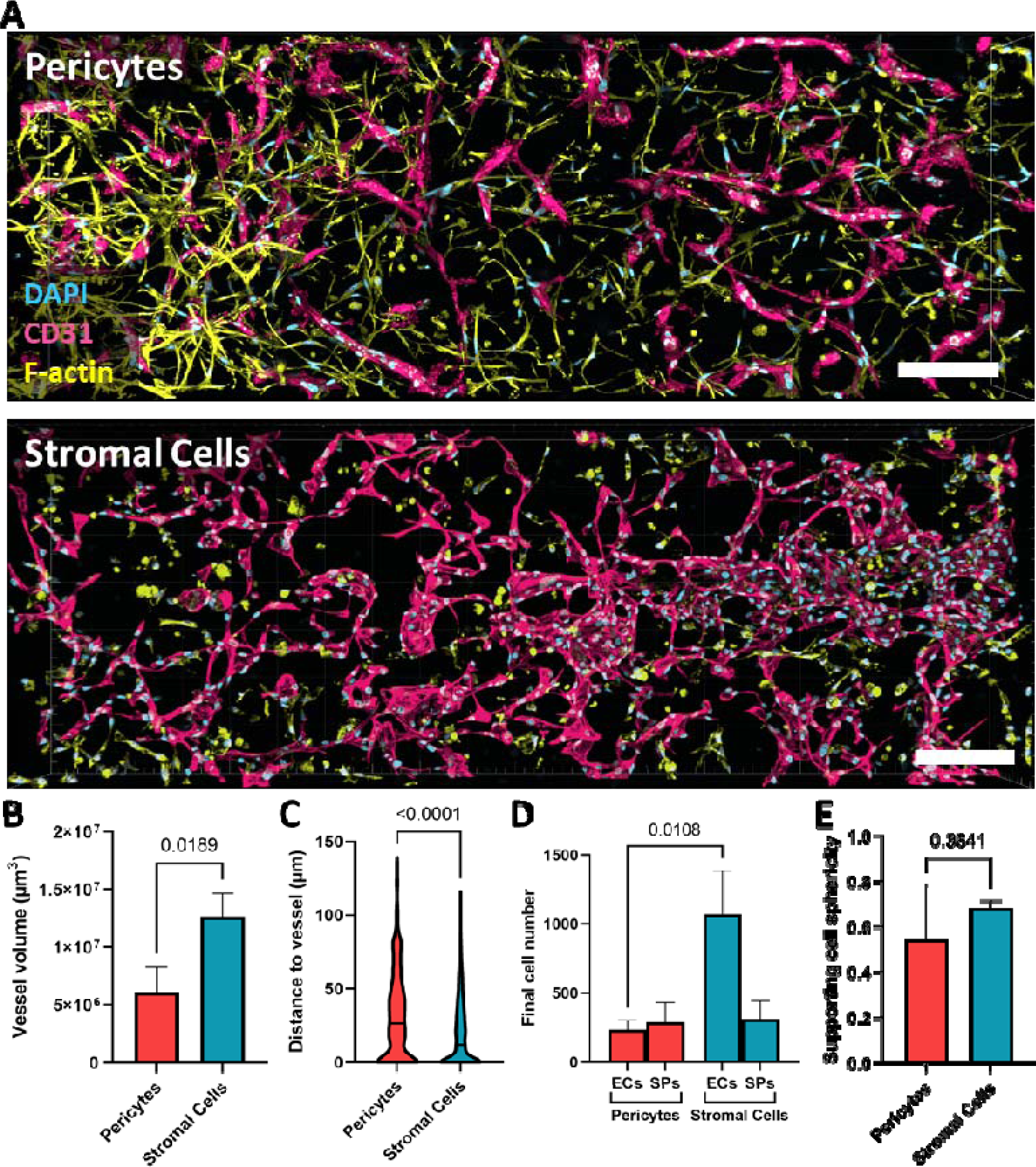
Comparison of pericytes and stromal cells in promoting vasculogenesis of endothelial cells. **(A)** Immunofluorescence images of the vasculogenesis process within the microfluidic chips co-cultured with either pericytes or stromal cells, stained for nuclei (DAPI; cyan), CD31 showing the endothelial vessel-like structures (magenta) and the supporting cells’ actin cytoskeleton (Phalloidin; yellow). Scale bars: 200 μm. **(B)** Comparison of total vessel volume between both co-culture conditions (n = 3, unpaired t-test, *p*<0.05 is considered significant). **(C)** Comparison of the distance between each type of supporting cell and the nearest vessel-like structure. The trunked violin plots depict summary statistics and the kernel density estimation to show the frequency distribution of each condition. The middle line represents the median (n = 3, Mann-Whitney test, *p*<0.05 is considered significant). **(D)** Comparison of the final total number of endothelial cells (ECs) and supporting cells (SPs) per imaged section, in the two conditions (n = 3, unpaired t-test, *p*<0.05 is considered significant). **(E)** Comparison of supporting cell’s average sphericity (n = 3, unpaired t-test, p<0.05 is considered significant).

In co-culture with endothelial cells, the stromal cells seem to have lost the elongated phenotype after the ten days of culture, a phenomenon that cannot be found in the angiogenesis setup. A major subset of the stromal cells are the fibroblasts, a cell type highly specialized in matrix degradation and deposition (Viswanathan *et al*., 2019). In wound healing, fibroblasts release a great amount of proteases that degrade fibrin networks (Plikus *et al*., 2021). During the remodeling, especially in low-density hydrogels like the one used in this study, fibroblasts could temporarily lose their attaching points and become more rounded (Fisher, Varani and Voorhees, 2008; Tamariz and Grinnell, 2002). However, a comparison of the supporting cell type morphology was performed based on average sphericity and no significant difference was found, partially due to the donor-to-donor variability (Figure 3E).

Overall, in our study, the endothelial structures formed in the presence of the stromal cells resemble the bloated and disorganized networks of the primitive plexus typical of the vasculogenesis processes that occur in the early embryo (Udan, Culver and Dickinson, 2013). In contrast, pericytes seem to inhibit this process and promote the formation of structures more similar to the vessels found in later stages of development. At the same time, pericytes seem to arrest the proliferative capacity of the endothelial cells, leading to a smaller final endothelial volume.

The main identifier of pericytes *in vivo* is the presence of a dense fiber network shared with the endothelial vessels, known as the basement membrane (Jayadev and Sherwood, 2017). Confirming the previous findings, transmission electron microscopy (TEM) reveals a close association of the pericytes with the endothelial vessels, forming a common basement membrane of shared matrix fibers (Figure 4A). Such an interaction is rarely observed between the stromal cells and the endothelial cells, which only present a loose association without a remarkable amount of shared fibers (Figure 4B). The incubation of the samples with tannic acid, which stains collagen and elastin in black, reveals that the composition of both matrixes is also different. The space between stromal and endothelial cells is filled with dark fibrous collagen and elastin, while the basement membrane remains mostly unstained. The basement membrane is a combination of laminin, collagen IV, fibronectin, and many other proteins, explaining the gray tone of the interface (Jayadev and Sherwood, 2017). The basement membrane is interrupted by contact zones between the pericyte and the endothelial cell membranes (Figure 4A, arrows), allowing direct cell-cell communication through integrins and other transmembrane molecules (Picoli, Birbrair and Li, 2024). The areas where the endothelial cells attach to themselves or others by tight junctions, such as Claudin-5, or by adherens junctions, such as CD31 and VE-Cadherin, are also visible in the TEM images (Figure 4A and B, asterisks)(Dejana and Orsenigo, 2013).

**Figure 4.**
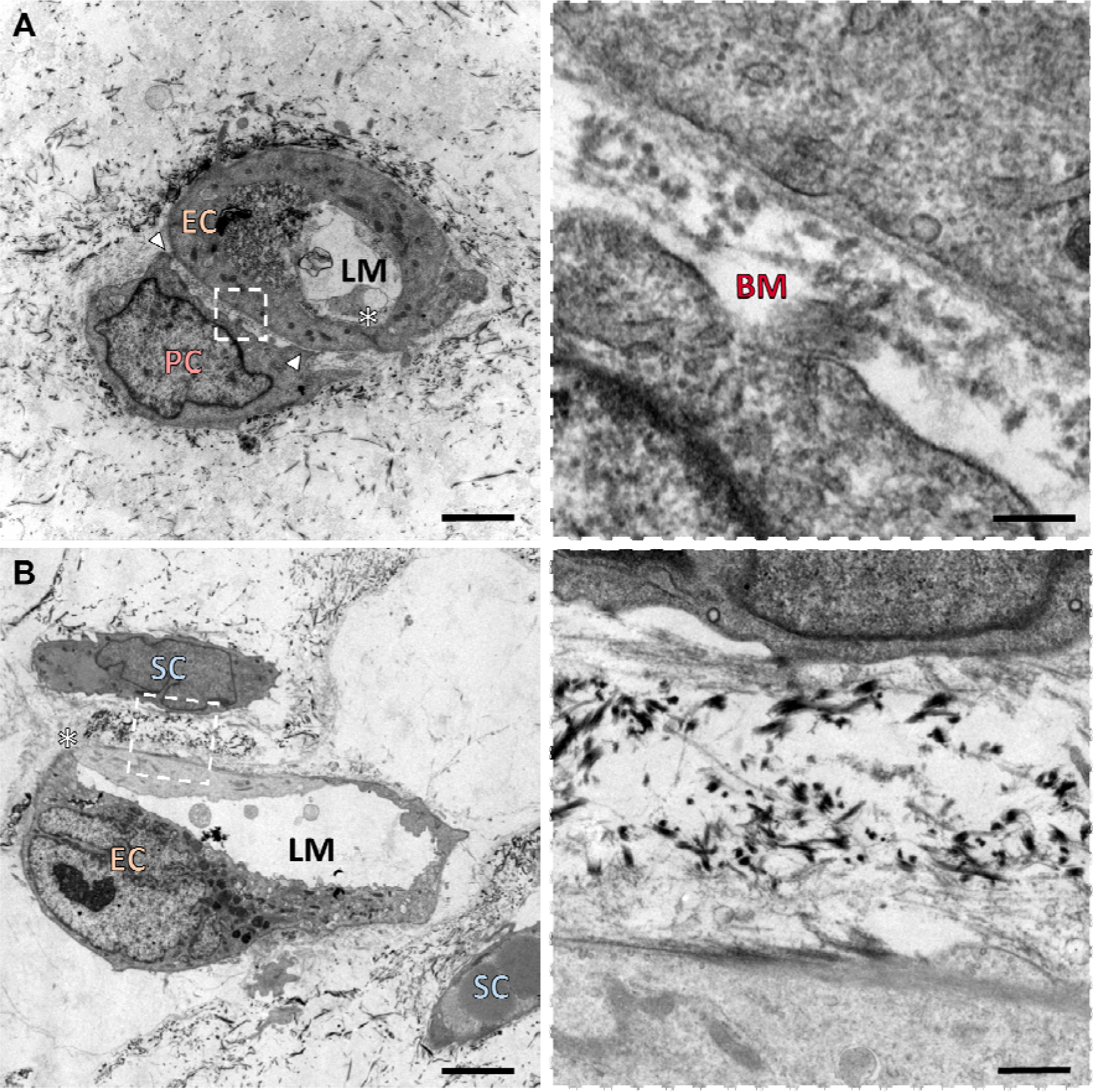
TEM image of the contact between supporting cells and vessels (**A**: pericytes, **B**: stromal cells). Only pericytes possess a common basement membrane matrix with the endothelial cells. EC: Endothelial Cell; PC: Pericyte; SC: Stromal Cell; BM: Basement membrane; LM: Lumen; Arrows: pericyte-endothelial cell junction; *: endothelial junction. Scale bars: 2500 nm (top left), 5000 nm (bottom left), 250 nm (right).

### 2.4. A transcriptomics insight into supporting cells differential behavior

We used RNA-seq to comprehend the differences in vasculogenesis in how pericytes and stromal cells regulate endothelial cell division and organization differently. Briefly, vasculogenesis gels with endothelial-supporting cell-co-cultures incubated for ten days were minced and dissolved to isolate RNA and perform bulk 3’mRNA-sequencing. Heatmap and dendrogram (Figure 5A) show the 1,550 significantly differently expressed genes (signature genes) between the different supporting cell types (adjusted p-value p<0.05). These differently expressed genes are evenly distributed between both cell types (Figure 5B). To better visualize the most relevant results, the 200 signature genes of each co-culture condition with the highest fold change were plotted in a network (Figure 5C) showing the relations (edges) between genes (nodes), removing the genes with no known interaction.

**Figure 5.**
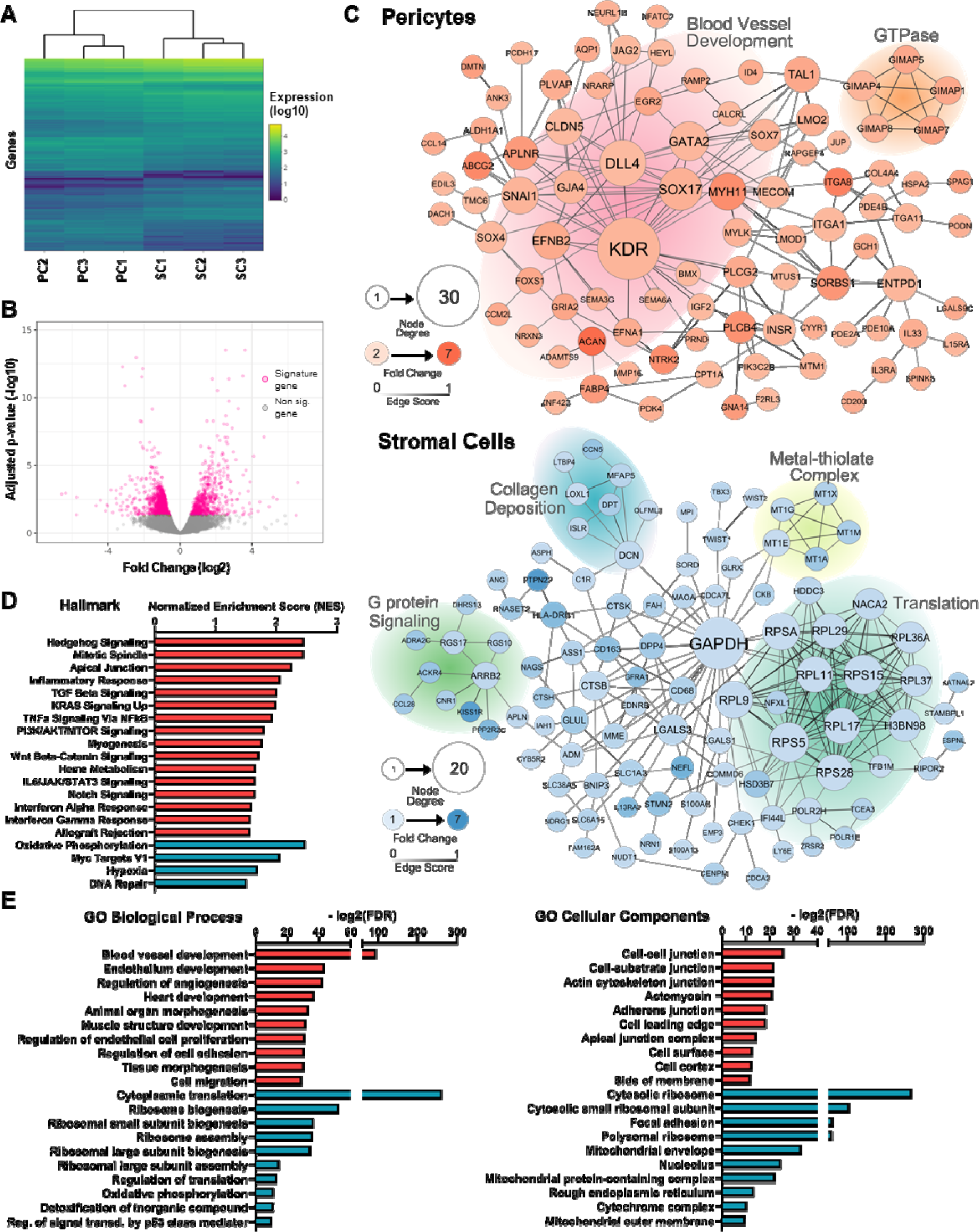
Gene expression analysis of pericyte and stromal cell co-cultures with endothelial cells. **(A)** Dendrogram and heat map of hierarchical clustering of the 1,550 differently expressed genes between the three pericyte (PC) and stromal cell (SC) donors (n = 3, for further explanations in the statistical analysis see the Methods section). **(B)** Volcano plot of the 1,550 differently expressed genes plotting their fold change (log2) by adjusted p-value (-log10). Positive fold change values are signature genes in pericytes, while negative ones are in the stromal cells. **(C)** Network visualization of the top differently expressed genes in both types of co-cultures. Each node represents a signature gene and the connections to other genes show a known relation in their expressions. The node color visualizes the fold change of its expression, and the size represents its importance in the network, understood as the number of relations with other genes. The tone of the connections shows the “Edge score”, the strength of the relation between them. The background-shading groups are merely orientative and represent relevant gene sets based on the GO databases. **(D)** Hallmark gene sets from MsigDB on the differentially expressed genes between pericytes (red) and stromal cell (blue) co-cultures. The scale bar represents the Normalized Enrichment Score (NES). **(E)** Ontological analysis of the differently expressed genes based on GO databases, of pericytes (red) and stromal cells (blue). The scale bar represents the adjusted False Discovery Ratio (FDR).

Gene set enrichment analysis (GSEA) of all signature genes shows that pericytes promote the upregulation of genes related to blood vessel formation and organization (Figure 5D and E). No known signaling gene is exclusive of either adult angiogenesis or vasculogenesis, since both involve activation of the same pathways (Patel-Hett and D’Amore, 2011). Within them, Notch overexpression in the pericyte co-culture is central in the regulation of EC behavior during neovascularization (Fernández-Chacón *et al*., 2021). The detection of free VEGFA by the KDR/VEGFR2 receptor activates an EC as a “tip cell”, suppressing its proliferation and driving it to lead vessel elongation. Delta-like ligands (DLL4) on tip cells interact with Notch receptors (Notch1 and Notch4) on adjacent cells, inhibiting their tip cell behavior and marking them as “stalk cells” (Hellström *et al*., 2007). This process is crucial to control tubulogenesis and avoid aberrant networks.

Elaborating on other upregulated pathways in pericytes according to the Hallmark analysis, Sonic Hedgehog (SHH) coordinates both embryological and post-natal vascularization through both its canonical and non-canonical pathways, promoting EC survival, differentiation, and maturation (Salybekov *et al*., 2018). SHH upregulates Notch1 in both endothelial cells and pericytes. The Wnt pathway, on the other hand, regulates vascularization through β-catenin-dependent mechanisms that affect cytoskeleton organization, directly affecting vessel tubulogenesis, remodeling, and stability (Olsen *et al*., 2017). The formation of vessel lumen is dependent on mitotic spindle orientation since it will establish the apical-basal polarity of the endothelial cells (Wu, Zhou and Li, 2020). The apex of the lateral membrane of these adjacent polarized endothelial cells is joined by apical junctions, which have a major role in regulating cell-cell adhesion and paracellular permeability (Rusu and Georgiou, 2020; Wang and Margolis, 2007).

The transforming growth factor beta (TGF-β) pathway regulates pericyte attraction and activity and acts as a cytostatic by the inhibition of c-Myc expression, stabilizing new vessels (Guerrero and McCarty, 2017). Both phosphoinositide 3-kinase-AKT-mammalian target of rapamycin (PI3K-AKT-mTOR) and Kirsten rat sarcoma virus (KRAS) are related pathways activated in endothelial cells by pro-angiogenic molecules and act regulating the secretion of VEGF, nitric oxide and Ang1/2 (Mendoza, Er and Blenis, 2011).

The RNAseq analysis shows an upregulation of the pro-inflammatory interleukin-6 (IL-6) in pericytes with respect to unselected stromal cells. The IL-6 receptor is highly expressed in pericytes, and its binding activates STAT3, increasing mural cell recruitment and the release of pro-vasculogenic growth factors (Didion, 2017; Villar-Fincheira *et al*., 2021). Other pro-inflammatory cytokines such as tumor necrosis factor-alpha (TNFa), interferon alpha and gamma, and the NF-kB signaling pathways are also downregulated in stromal cells. Stromal cells are widely known to have immunomodulatory properties which are being explored in cell therapy (Bourin *et al*., 2013; Caplan, 2017; Galderisi, Peluso and Di Bernardo, 2022; Melief *et al*., 2013; Viswanathan *et al*., 2019). This is in line with the downregulation of many inflammatory molecules observed in that co-culture condition (Patrikoski, Mannerström and Miettinen, 2019).

The GO Cellular Component analysis reveals overexpression of cell-cell and cell-matrix junctions in the pericyte co-cultures, such as Claudin 5 (CLD5) and many types of integrins (ITGA) (Dejana and Orsenigo, 2013; Mongiat *et al*., 2016). The tightness of the endothelial barrier is dependent on these cellular connections, essential to maintain the structural integrity of the vessels and avoid the leakage of the blood contents. Some of the upregulated integrins are also directly involved in vasculogenesis such as ITGA1 or 11 (Mongiat *et al*., 2016).

Stromal cell co-culture shows a robust upregulation of Myc targets and hypoxia pathways, both profoundly involved in cell proliferation (Batie *et al*., 2019; Bretones, Delgado and León, 2015; Kaluz, Kaluzová and Stanbridge, 2008; Pugh and Ratcliffe, 2003). Other pro-mitotic signaling such as p53 and G protein cascades are also upregulated (Carlton, Jones and Eggert, 2020; Engeland, 2022). The consequences are shown by the increased cytoplasmic translation and ribosomal, mitochondrial, and nucleolar biogenesis, intrinsic parts of cell division (Carlton, Jones and Eggert, 2020). The increased DNA repair mechanisms and oxidative phosphorylation may indicate a response to the increased metabolic activity and replication stress associated with cell proliferation (Magdalou *et al*., 2014; Saxena and Zou, 2022). Stromal cells also overexpress genes involved in collagen genesis and deposition, explained by the large percentage of fibroblasts present in this population (Sun *et al*., 2019).

Not all differences in gene expression can be translated to protein amount and signaling since RNAseq is blind to post-transcriptional regulation and protein stability and degradation. However, there is a very clear trend of upregulated vasculogenesis and tubulogenesis genes in the co-cultures supported by pericytes, including Notch, TGF-β, IL-6, SHH, and Wnt. On the other hand, stromal cells lead to the overexpression of genes related to cell division and organelle biogenesis (such as Myc and p53), together with the associated cell stress and accelerated metabolism; and the downregulation of inflammatory cytokines that characterize this population. This is in line with the previously shown morphological and immunofluorescence results (Figures 3 and 4).

## 3. Conclusion

The relationship between pericytes and mesenchymal stromal cells has been widely debated in the last years, ranging from considering them as two completely independent populations to suggesting the latter as an activated form of the pericyte (Blocki *et al*., 2013; Blocki *et al*., 2018; Guimarães-Camboa *et al*., 2017; Souza *et al*., 2016). It has also been discussed if the pericytic identity can be maintained in culture or if subsequent passages outside of the body would lead to a loss of its characteristics (Blocki *et al*., 2013).

Here, we demonstrate that the isolation of pericytes from the stromal vascular fraction of the adipose tissue based on the expression of the MCAM marker provides a population of plastic-adherent cells that conserves their ability to act as mural cells *in vitro* for at least five passages. Both immunofluorescence and TEM imaging proved this interaction. We also demonstrate that these pericytes act significantly differently from their unselected counterpart, promoting longer and more branched sproutings in angiogenesis. In vasculogenesis, on the other hand, pericytes arrest endothelial cell division and stimulate their arrangement in vessel-like structures, while stromal cells stimulate endothelial cell proliferation, forming aggregates similar to the primitive capillary plexus. This is also reflected in the transcriptomics analysis, which shows that pericytes upregulate pathways related to vessel formation, such as VEGFA-VEGFR-DLL4-Jagged1-Notch, SHH, Wnt, IL6-JAK-STAT3 and PI3K-AKT-mTor; while stromal cells stimulate cell proliferation through pathways associated to Myc and p53 and decrease pro-inflammatory signaling.

These results are relevant both to understand the function of pericytes and stromal cells *in vitro* and to improve current strategies for tissue-engineered vascularization. For instance, the propensity of stromal cells to induce endothelial proliferation could be harnessed to support the first steps of vasculogenesis, while the vessel-stabilizing influence of pericytes might be critical in preventing pathological vascularization.

## 4. Experimental Section

### 4.1. Cell isolation and culture

All human tissues were obtained following the Declaration of Helsinki after informed consent at RWTH Aachen University Hospital, Aachen, Germany. Stromal cells and pericytes were isolated from adipose tissue kindly provided by the Clinic for Plastic, Hand and Burns Surgery (RWTH Aachen University Hospital, ethics approval reference number EK197-19). First, the adipose layer was separated from the dermis and manually minced until achieving a homogenous slurry. The minced fat was added to gentleMACS C-Tubes (Miltenyi), mixed with the same volume of collagenase I (1 mg/mL; Sigma-Aldrich), and placed in a gentleMACS Octo Dissociator to achieve complete homogenization and digestion at 37°C with consecutive stirring steps for 15 minutes. The mixture was left for another 30 minutes at 37°C in soft agitation on a roller mixer. The digested SVF was then filtered using a 100 μm cell strainer (Falcon), mixed with sterile PBS, and centrifuged at 300 g. The supernatant was removed and the pellet was resuspended in PBS for two more washing steps. To expand the stromal cells, one-quarter of the SVF suspension was seeded in a cell culture-plastic flask and cultured in Mesenpan media (PAN-Biotech) supplemented with 2% (v/v) fetal bovine serum (FBS; Gibco) and 1% (v/v) antibiotic/antimycotic solution (ABM; Gibco).

The other part of the SVF is diluted 1:10 in Erythrocyte Lysis Buffer (PAN-Biotech), briefly vortexed, and incubated at room temperature (RT) for 10 minutes. The mixture was washed another 3 times as previously described. The resulting pellet was resuspended in 10 μL anti-CD146 magnetic microbeads (Miltenyi) and 90 μL blocking buffer (Miltenyi), and incubated for 15 minutes at 4°C. The mixture was washed again three times to remove unattached magnetic microbeads. A MACS LS column (Miltenyi) was attached to a magnetic separator (Miltenyi) and prepared by rinsing it with PBS. The labeled cell suspension was added to the column and rinsed three times with Mesenpan to remove the CD146^-^ cells. The column was then separated from the magnetic holder, filled with media, and pressed with a sterile plunger. The resulting CD146^+^ cell suspension was considered as pericytes and cultured as previously described in the same conditions as the stromal cells.

Umbilical cords were kindly provided by the Centralized Biomaterial Bank of the RWTH Aachen University (cBMB) and the Department of Gynecology and Perinatal Medicine according to RWTH Aachen University, Medical Faculty Ethics Committee (cBMB project number 323). Human umbilical vein endothelial cells (HUVECs) were isolated from umbilical cords using collagenase type I (Sigma-Aldrich, 400 U/mL) to separate the cell layer surrounding the vein lumen. The detached cells were subsequently expanded in endothelial cell growth medium 2 (EGM2; Promocell) supplemented with 1% (v/v) ABM. The human umbilical cords were generously provided by the Clinic for Gynecology and Obstetrics (RWTH Aachen University Hospital) and cells were isolated within 24 hours after delivery.

HUVECs, stromal cells, and pericytes were cryopreserved in liquid nitrogen using a freezing solution consisting of 80% (v/v) DMEM, 10% (v/v) FBS, and 10% (v/v) DMSO. Prior to the device experiment, HUVECs were thawed and expanded up to passage 3 in EGM2. Stromal cells and pericytes were cultured up to passage 4 in Mesenpan.

### 4.2. Preparation of the vascularization assays in microfluidic devices

HUVECs, pericytes, and stromal cells were detached by trypsin/EDTA (Pan-Biotech) digestion. Cells were then suspended in tris-buffered saline (TBS) solution containing thrombin (Sigma-Aldrich) and calcium chloride (CaCl_2_; Sigma-Aldrich). This mixture was combined with a fibrinogen solution (VWR) pre-cooled on ice and either 10 μL were injected into the central channel of an idenTx 3 microfluidic chip (AIM Biotech) for the chip experiments or 350 μL were directly deposited into 24-well plate wells for the bulk gels. The resulting gel contained 5 mg/mL fibrinogen, 3 IU/mL thrombin and 3.75 mM CaCl_2_.

For imaging after the preparation, cells were pre-stained with Vybrant dyes (Invitrogen) by adding 5 μL of cell-labeling solution to a 1×10^6^ cell/mL cell suspension in serum-free cell media. The mixture was vortexed and incubated for 20 minutes at 37°C. Then, cells were centrifuged at 300 g and washed with PBS three times to remove residual dye.

For the vasculogenesis chip set-up, 10×10^6^ HUVECs/mL and 2×10^6^ supporting cells/mL were pre-mixed in the injected solution. For the bulk vasculogenesis gel, 10×10^6^ HUVECs/mL and 5×10^6^ supporting cells/mL were added. For the angiogenesis chip set-up, only 2×10^6^ supporting cells/mL were suspended in the polymerizing solution. In all cases, the fibrin gels were incubated for 20 min at room temperature and another 20 min at 37°C and 5% CO_2_ in a humidified atmosphere to achieve complete polymerization of the fibrin. For the angiogenesis assay, the bottom channel was filled with a solution of 50 μg/mL fibronectin in EGM microvascular vessel 2 (EGM-MV 2, PromoCell) immediately after the polymerization and incubated for 40 minutes at 37°C. The channel was washed with EGM-MV2 and a solution of 2×10^5^ HUVECs/mL was injected in the coated channel.

The reservoirs of all chips were filled with EGM-MV 2 supplemented with 1% (v/v) ABM, adding 100 μL to the bottom left well, 80 μL to the bottom right well, 60 μL to the top left well and 40 μL to the top right well, to achieve gravity-driven flow both inside each of the channels and between them through the hydrogel. The chips were placed in humidified chambers with the bottom exposed to air, in a 5% CO_2_ and 37°C atmosphere. The media was changed every 24 hours for 10 days. For the bulk gels in 24-well plates, 500 μL of EGM-MV 2 was added and changed every 48-72 hours. Three biological replicates (independent donors) were used.

For fixation, the medium was removed from the channels and substituted with ice-cold methanol at -20°C. All fibrin gels were incubated at 4°C for 30 minutes and then washed three times with PBS.

### 4.3. Immunofluorescence imaging

The fixed chips were first stained with mouse anti-CD31 (PECAM-1, 1:100; Sigma-Aldrich) solved in antibody diluent made of 3% Bovine Serum Albumin (BSA) and 1% ABM for 48 hours at 37°C. Then, the samples were washed three times. Afterward, a mixture of Alexa Fluor® 594 goat anti-mouse IgG (1:400, Invitrogen) and Phalloidin-iFluor™ 488 Conjugate (Cayman Chemical) was added to the samples and incubated for 48 hours at 37°C. The chips were washed, stained with 0.4 μg/mL of a DAPI solution for 2 hours at 37°C, washed again, and stored at 4°C until imaging. Microscopy was done with a LSM 980 with Airyscan 2 (Zeiss) confocal laser scanning microscope, creating stacks of the whole gel thickness, 200 μm, with a step size of 3 μm. The resulting composite was processed with Imaris 10 (Oxford Instruments), creating 3D surfaces of the CD31^+^ vessel-like objects. The volume of these objects was measured. The measure function of ImageJ was used to calculate the angiogenic sprouting length. The sproutings were considered tubular and homogenous enough to assess them as cylinders, approximating thir diameter based on the volume and lenght data. The number of branching points was manually counted. The vessel-like objects were subtracted from the F-actin channel to show only the supporting cells. From this, 3D surfaces were again created to measure the distance of these cells from the CD31^+^ structures.

Normal distribution of the data was checked using the Shapiro-Wilk test. Except for the “distance to vessel” data of the angiogenesis assay, all data were normally distributed and unpaired t-tests were performed. For the “distance-to-vessel” data, the non-parametric Mann-Whitney test was used.

### 4.4. TEM imaging

The bulk fibrin gels were fixed in a 3% glutaraldehyde solution for at least 24 hours. They were then rinsed in 0.1 M Soerensen’s phosphate buffer (Merck) and fixed with 1% osmium tetraoxide (Carol Roth) in a 25 mM sucrose buffer (Merck). After a dehydration process with a series of increasing ethanol concentrations, the samples were immersed in propylene oxide, then in a 1:1 mixture of Epon resin (Serva) and propylene oxide for one hour, followed by a final one-hour incubation in pure Epon resin. The samples were then embedded in pure Epon at 90°C for two hours.

Ultrathin sections between 90 to 100 nm were obtained using an ultramicrotome (Reichert Ultracut S, Leica) equipped with a Diatome diamond knife. These sections were placed onto Cu/Rh grids (HR23 Maxtaform, Plano). To enhance contrast, the sections were stained with a 0.5% uranyl acetate and 1% lead citrate solution (both from EMS). The final observation was performed using a Zeiss Leo 906 (Carl Zeiss) transmission electron microscope, operated at an acceleration voltage of 60 kV.

### 4.5. RNA isolation, sequencing and analysis

RNA isolation was performed using TRIzol (Invitrogen) and following the manufacturer’s user guide. Briefly, cold TRIzol was added to the fibrin gel and incubated for 5 minutes. The mixture was then pipetted up and down until the gel disagreggated, and stored at -20 C until isolation. After thawing, chloroform was added to the sample, incubated for 3 minutes, and centrifuged for 15 minutes at 12,000 g at 4 C to separate the phases. The top aqueous phase containing the RNA was carefully extracted and placed in a different tube. Then, isopropanol was added and the tubes were centrifuged again to precipitate the RNA. The supernatant was discarded, RNA pellet was resuspended in 75% ethanol and centrifuged again. This process was repeated three times to remove isopropanol leftovers from the mixture. Finally, the RNA pellet was air-dried until no ethanol was left and resuspended in RNase-free water.

3’mRNA-Seq libraries were prepared using Lexogen QuantSeq 3’mRNA-Seq v2 Library Prep Kit FWD with UDIs following the manufacturer’s protocol. Prior to library preparation, the concentration of RNA was measured using the Promega Quantus Fluorometer. Additionally, the size distribution of the RNA was assessed using the Agilent TapeStation with an RNA ScreenTape. Quantification and quality assessment were repeated after library preparation, again using the Quantus fluorometer and the Agilent TapeStation with a High Sensitivity D1000 ScreenTape. Libraries were denatured, diluted, and loaded onto a NextSeq High Output v2.5 (75 cycles) flow cell. The 1% PhiX control library was spiked in to improve base calling accuracy. Single-end sequencing was performed with 75 cycles on the Illumina NextSeq platform according to the manufacturer’s instructions.

FASTQ files were generated using bcl2fastq (Illumina). To facilitate reproducible analysis, samples were processed using the publicly available nf-core/RNA-seq pipeline version 3.12 (Ewels *et al*., 2020) implemented in Nextflow 23.10.0 (Di Tommaso *et al*., 2017) with minimal command. In brief, lane-level reads were trimmed using Trim Galore 0.6.7 (Felix Krueger *et al*., 2023) and aligned to the human genome (GRCh39) using STAR 2.7.9a (Dobin *et al*., 2013). Gene-level and transcript-level quantification was done by Salmon v1.10.1 (Patro *et al*., 2017). Heatmap/dendrogram and volcano plot visualizations were performed using custom scripts in R version 4.3.2 using the DESeq2 v.1.32.0 framework (Love, Huber and Anders, 2014).

For the hallmark gene set analysis, Gene Set Enrichment Analysis (GSEA) was performed using clusterProfiler version 4.6.2. Pre-ranked gene lists, generated from differential gene expression analysis, were input into clusterProfiler to identify enriched biological pathways (Liberzon *et al*., 2015). Enriched hallmark gene sets with an FDR q-value < 0.05 were considered significant.

Gene network visualization and functional enrichment of GO pathways were done using the stringApp v2.0.3 (Doncheva *et al*., 2019) of CytoScape 3.10.1 (Shannon *et al*., 2003). For the gene networks, the top 200 genes with higher adjusted p-values of each co-culture type were plotted using the STRING visualization. Singletons and pairs were removed, and the remaining network was organized with the Prefuse Force Directed Layout. Then, enrichment data for all the signature genes of GO Biological Processes and GO Cellular Components was exported with a redundancy cutoff of “0.7” to remove nearly identical terms. The ten pathways with the highest adjusted FDR of each condition were plotted using GraphPad Prism 9.5.1.

## Acknowledgements

We want to thank the Department of Gynecology and Perinatal Medicine (Univ.-Prof. Dr. Stickeler) and the Centralized Biomaterial Bank of the RWTH Aachen University (cBMB) for providing the human umbilical cords. We also wish to express our gratitude to the Clinic for Plastic, Hand and Burns Surgery (Univ.-Prof. Dr. Beier) for providing fat tissue (University Hospital Aachen, RWTH University Aachen, Germany). We thank Hana Masutani for helping with the illustrations in Figure 1A and B. This work was supported by the Confocal Microscopy Facility and the Genomics Facility of the Interdisciplinary Center for Clinical Research (IZKF) Aachen, within the Faculty of Medicine at RWTH Aachen University. We thank the Electron Microscope Facility (Institute of Pathology, RWTH Aachen University Hospital) for their support in the TEM, and particularly to Dr. Eva Miriam Buhl for her advices in image interpretation.

